# The protist ubiquitin ligase effector PbE3-2 targets RD21A to impede plant immunity

**DOI:** 10.1101/2022.10.26.513834

**Authors:** Chao Li, Shaofeng Luo, Lu Feng, Qianqian Wang, Jiasen Cheng, Jiatao Xie, Yang Lin, Yanping Fu, Daohong Jiang, Tao Chen

## Abstract

Clubroot caused by the soil-borne protist pathogen *Plasmodiophora brassicae* is one of the most devastating diseases of Brassica oil and vegetable crops worldwide. Understanding the pathogen infection strategy is crucial for the development of disease control. However, the molecular mechanism by which this pathogen promotes infection remains largely unknown. Here, we identified a *P. brassicae*-secreted effector PbE3-2 that impedes plant immunity by ubiquitinating the immune regulator RD21A for degradation. Overexpression of *PbE3-2* in *Arabidopsis thaliana* resulted in higher susceptibility to *P. brassicae* and decreases in chitin-triggered reactive oxygen species burst and expression of marker genes in salicylic acid signaling. PbE3-2 interacted with and ubiquitinated RD21A *in vitro* and *in vivo*. Mutant plants deficient in *RD21A* exhibited similar susceptibility and compromsied immune responses as in *PbE3-2* overexpression plants. These results suggest that *P. brassicae* promotes clubroot disease through RD21A degradation mediated by the effector PbE3-2. As PbE3-2 is widely conserved across different *P. brassicae* pathotypes, the degradation of RD21A by PbE3-2 might be a prevalent infection strategy in this pathogen.

## Introduction

*Plasmodiophora brassicae* is an obligate biotrophic pathogen and the causal agent for clubroot disease worldwide. It parasitizes the roots of cruciferous plants, resulting in the formation of club-shaped galls. The above-ground part of the plant tends to show the symptoms of wilting, stunting, yellowing, premature senescence or death (Hwang et al, 2012). Clubroot disease causes serious damage to crop yield and quality, resulting in a global loss of 10%–15% in yield and 2%–6% in oil content (Saharan *et al*, 2021). This disease is now found in all Brassica-growing areas of the world and seems to be getting worse everywhere, which is notoriously known as “the killer” of rapeseed. Breeding and the cultivation of resistant varieties are the most effective approach to mitigate the threats of this disease (Guo *et al*, 2022). However, new virulent pathotypes of *P. brassicae* have constantly emerged (Galindo-Gonzalez et al, 2020). Understanding how *P. brassicae* successfully infect plants will facilitate the breeding of new resistant cultivars and the development of disease control.

*P. brassica*e is an obligate biotrophic protist belonging to a subgroup of Rhizaria, one of the most poorly understood subgroups of eukaryotes. The obligate lifestyle limited to investigating molecular mechanisms of pathogenic infection. Many eukaryotic biotrophic plant pathogens have evolved advanced strategies to deliver the effector proteins into the host cell during infection to manipulate plant defenses and facilitate parasitic colonization (Dodds & Rathjen 2010). Identification of effectors of *P. brassicae* is critical for understanding how the pathogen manipulates the immune response of plants and regulates clubroot resistance. The availability of the genome and transcriptome of *P. brassicae* pathotypes provides an opportunity for identifying the putative effectors (Schwelm A. et al, 2015a; Bi et al, 2016; Rolfe *et al*, 2016). *P. brassicae* can secrete effectors into host cells to cause extensive cellular and tissue rearrangements (Ludwig-Müller *et al*, 2015). *P brassiae* effector SSPbP53 can inhibit cruciferous papain-like cysteine proteases to suppress plant immunity (Perez-Lopez et al, 2021). PBZF1 is an RxLR effector interacting with host kinase SnRK1.1, and inhibits the biological function of SnRK1.1. Heterologous expression of *PBZF1* in *A. thaliana* increased the susceptibility of plants to *P. brassicae* (Chen et al, 2021). To better understand the pathogenic mechanism of *P. brassicae*, it is necessary to further study the roles and molecular mechanisms of *P. brassicae* effector candidates.

Ubiquitination, which is an important post-translational modification of proteins unique to eukaryotes (Hershko & Ciechanover, 1998), serves as a degradation signal or changes the property of the target protein (Smalle and Vierstra, 2004). Usually, ubiquitin ligase E3 can specifically recognize target proteins and is therefore particularly important and extensively studied. Ubiquitination is involved in various mechanisms including cell cycle, apoptosis, protein degradation, and immune regulation, and is, therefore, one of the most important mechanisms regulating various life activities (Smalle and Vierstra, 2004; Hochstrasser et al, 2008; Kim et al, 2010; Ulrich & Walden, 2010). The *P. brassicae* e3 genome contains 139 putative E3 ubiquitin ligases, among which 115 harbor the conserved RING domain. Yeast invertase assay has verified that PbRING1 contains a functional signal peptide and has E3 ligase activity *in vitro* (Yu et al, 2019). However, the biological functions of E3 ubiquitin ligases in *P. brassicae* remain largely unknown.

Many studies have demonstrated that ubiquitination plays important roles in the regulation of plant immunity (Zhou & Zeng, 2017). *A. thaliana* E3 ubiquitin ligases PUB12/PUB13 and PUB22 ubiquitinate the pattern recognition receptor FLS2 and the exocyst complex subunit Exo70B2, respectively, to promote their degradation to attenuate PAMPs-triggered immune (PTI) signaling (Lu et al, 2011; Stegmann *et al*, 2012). Besides, PUB12/PUB13 also mediates the degradation of ABA co-receptor ABI1 to regulate ABA signaling (Kong *et al*, 2015). In addition, pathogens secrete effectors that can modulate the host ubiquitination system to influence plant immunity. The *Magnaporthe oryzae* effector AvrPiz-t targets the rice E3 ligases APIP6 and APIP10 forsuppressing PTI (Park *et al*, 2012; Park *et al*, 2016). The *Pseudomonas syringae* type III effector AvrptoB, which was shown to have E3 ubiquitin ligase activity, is important for the pathogenicity of *P. syringae* (Abramovitch et al, 2006). Understanding the functions of E3 ubiquitin ligases in *P. brassicae* can contribute to a better understanding of the molecular mechanism underlying the pathogenesis of *P. brassicae*.

This study aims to investigate the biological functions of secreted E3 ubiquitin ligases in *P. brassicae*, and understand the mechanisms for ubiquitination in clubroot development. Our findings suggest that the secreted E3 ubiquitin ligases ubiquitinate the plant cysteine protease RD21 thereby suppressing plant immunity for successful infection in protist pathogen.

## Results

### PbE3-2 is a RING type E3 ubiquitin ligase

By using Blast2GO to systemically investigate E3 ubiquitin ligases in the genome of *P. brassicae* ZJ-1 (Bi et al, 2019), we identified 60 genes encoding putative E3 ubiquitin ligases (Table EV1). We hypothesized that some of these E3 ubiquitin ligases are secreted into the host cell and contribute to successful infection. Five out of the 60 predicted E3 ubiquitin ligases possessed signal peptides, which were designated as PbE3-1, PbE3-2, PbE3-3, PbE3-4 and PbE3-5, respectively (Table EV1 and Appendix Fig. S1A). PbE3-1 and PbE3-5 were predicted to have transmembrane domain, while the remaining three PbE3s had no such domain (Appendix Fig. S1A). qPCR analysis indicated that these five PbE3s have differential expression patterns at different life stages of *P. brassicae* (Appendix Fig. S1B). PbE3-2 harbored a RING (Really Interesting New Gene) finger domain (Table EV1), and is conserved among *P. brassicae* pathotypes (Appendix Fig. S1C). The RING finger domain of PbE3-2 contained eight conserved Cys/His amino acid residues and thus belonged to the C_3_H_2_C_3_-type family (Fang & Weissman, 2004), and could coordinate two Zn^2+^ ions (Fig. 1A). The three-dimensional model of the RING finger domain in PbE3-2 was similar to that of the canonical RING (Fig. 1B). We first tested whether PbE3-2 has an enzymatic activity for self-ubiquitination. AtUBA1-S (E1 activating enzyme), AtUBC8-S (E2 conjugating enzyme), PbE3-2-Myc and His-FLAG-UBQ10 (ubiquitin) were expressed in *Escherichia coli* DE3 strains, and self-ubiquitination of PbE3-2 was tested in the presence or absence of E1 activating enzyme, E2 conjugating enzyme, and ubiquitin. PbE3-2 was converted into a mixture of high-M_r_ ubiquitinated protein products (Fig. 1B), suggesting that PbE3-2 is capable of self-ubiquitination. We also generated a PbE3-2-C423S mutant, in which the cysteine residue at position 423 in the conserved RING domain was substituted by serine. As a result, the C423S mutant protein showed no self-ubiquitination (Fig. 1C). Therefore, PbE3-2 is a E3 ubiquitin ligase.

**Figure 1.**
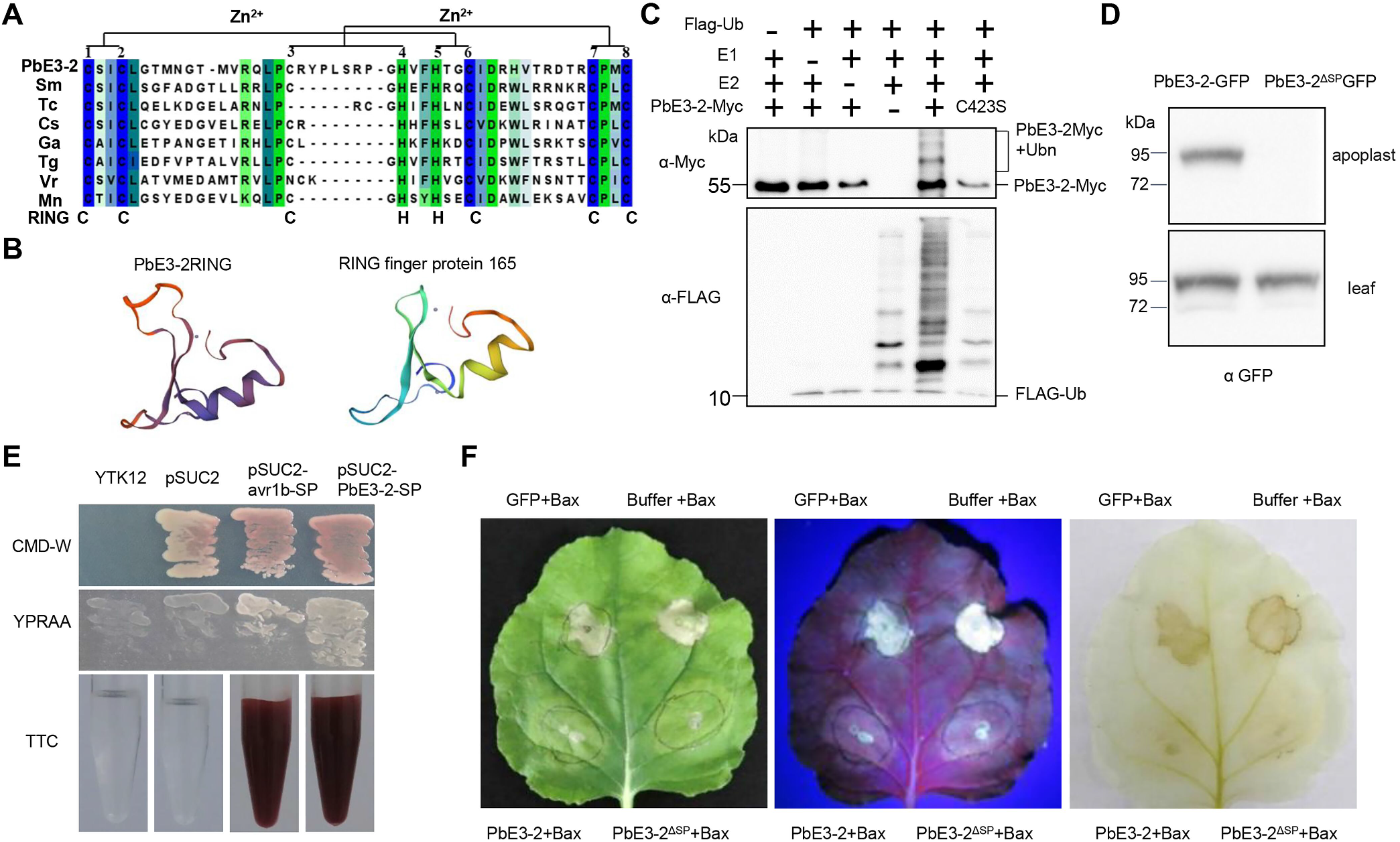
PbE3-2 is a typical secreted E3 ubiquitin ligase and suppressed BAX-induced cell death. A Protein alignment of PbE3-2RING and its homologs (*Symbiodinium microadriaticum* OLP82957.1, *Theobroma cacao* EOY12354.1, *Camelina sativa* XP_010413448.1, *Gossypium arboretum* KHG04456.1, *Toxoplasma gondii VEG* CEL71473.1’ *Vigna radiata var. radiata* XP_014517466.1, *Monoraphidium neglectum*, XP_013905359.1). B Structures of PbE3-2RING and RING finger protein 165 (*Zea mays*). C Self-ubiquitination of PbE3-2. Ub, E1, E2 and PbE3-2 are all present in the systems, PbE3-2 can undergo self-ubiquitination. The system lack of any component is a negative control. The PbE3-2 (C423S) mutant is also unable to undergo auto-ubiquitination. D PbE3-2-GFP and PbE3-2^ΔSP^ GFP infiltrated to *N. benthamiana* leaves for 2 days, apoplast washing fluid was extracted, and Western blotting with GFP antibody indicated that PbE3-2-GFP but not PbE3-2^ΔSP^ GFP was detected in the apoplast washing fluid. E PbE3-2 contains a functional signal peptide. Using a yeast signal-sequence trap system, culture filtrate of yeast expressing the plasmid pSUC2-SP PbE3-2, which expresses the PbE3-2 signal peptide under control of the pSUC2 promoter, produces a red color similar to that of the positive control of yeast expressing pSUC2-SPAvr1b, which contains the signal peptide from the *Phytophthora sojae* effector Avr1b. Yeast expressing the empty pSUC2 vector was used as a negative control. F PbE3-2 and PbE3-2^ΔSP^ suppressed BAX-induced cell death. PbE3-2 and BAX co-infiltrated to *N. benthamiana* leaves, BAX and PVX-GFP co-infiltrated as a negative control. Left picture was taken at 5 days after infiltration under visible light, middle picture was taken under UV light, right picture was decolorization with 75% acetic acid and 25% alcohol buffer. Experiments were independently performed three times with similar results.

### PbE3-2 suppresses BAX-induced cell death

As PbE3-2 harbors a putative signal peptide, to determine whether it is essential for protein secretion, we constructed two plasmids expressing PbE3-2-GFP fusion proteins with (PbE3-2-GFP) and without signal peptide (PbE3-2^ΔSP^-GFP), and delivered them into tobacco epidermal cells via *A. tumefaciens*. Apoplast washing fluid was extracted, and western blotting indicated that PbE3-2 but not PbE3-2^ΔSP^ was detected in the apoplast washing fluid (Fig. 1D), suggesting that PbE3-2 is a secreted protein. Yeast invertase assay showed that PbE3-2 harbors a functional signal peptide (Fig. 1E). These results are consistent with the notion that PbE3-2 is a secreted protein. Test of the ability to suppress BAX-induced cell death can help to identify functional effectors. Hence, PbE3-2 and BAX were co-infiltrated into the leaves of *N. benthamiana* using the *Agrobacterium-mediated* plant virus transient expression system (Fig. 2F). Both PbE3-2 and PbE3-2^ΔSP^ suppressed BAX triggered cell death in the infiltrated leaves, while the negative control GFP did not. These results suggested that PbE3-2 is an effector (Fig. 1D-F).

**Figure 2.**
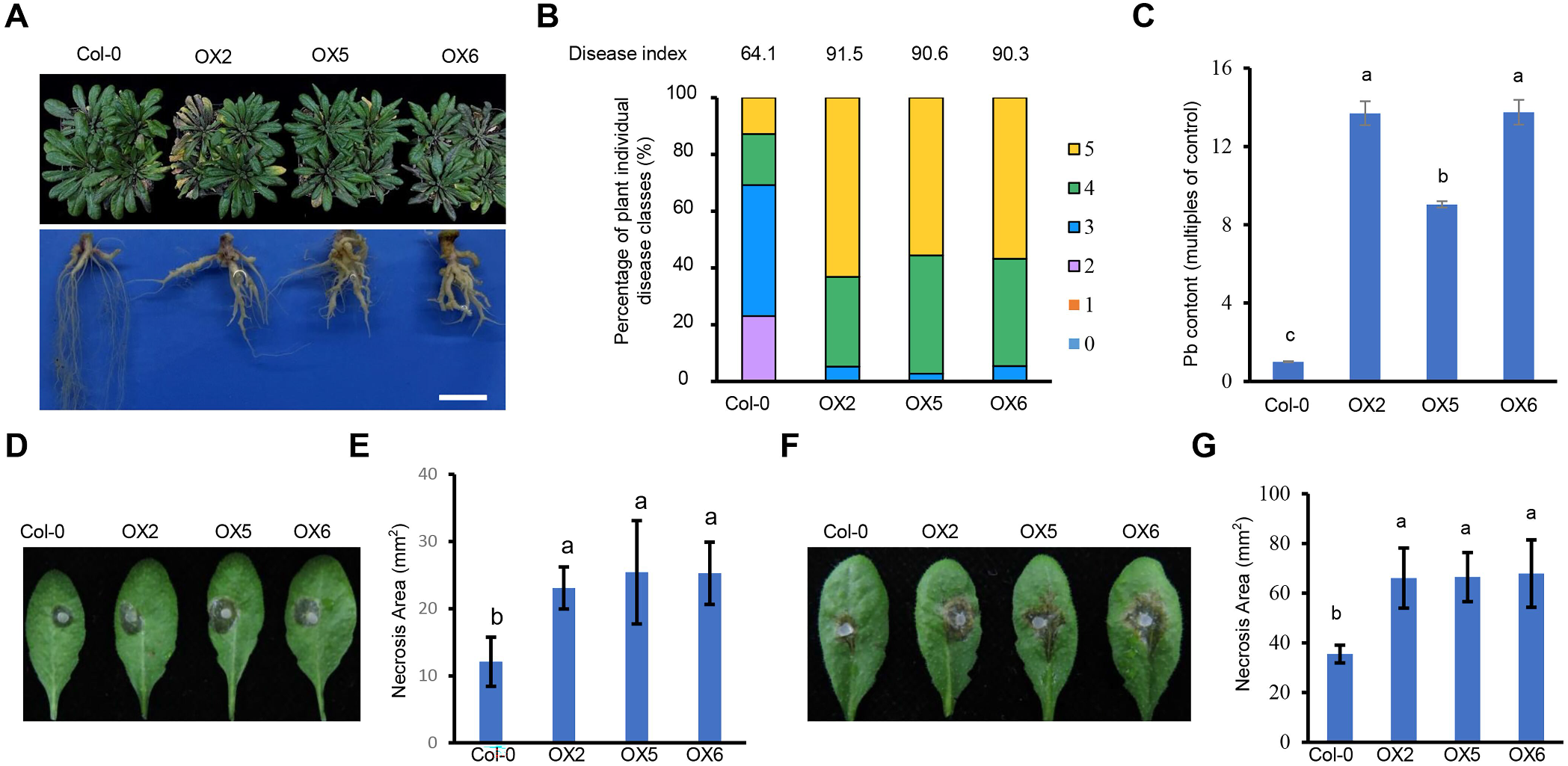
*PbE3-2* overexpression plants are more susceptible to pathogens. A Phenotype of Col-0 and *PbE3-2* overexpression plants inoculation with *P. brassicae* for 21 days. B Disease index of Col-0 and *PbE3-2* overexpression plants inoculation with *P. brassicae* for 21 days. Percentages of plants in the individual disease classes are shown. For transgenic lines and Col-0 plants, 35–40 plants were analyzed. The disease index for each sample is shown as a number above the respective histograms. The experiment was conducted with three independent replications. C Pathogen DNA quantification (Pb) by quantitative PCR, which is expressed as the percentage of the mean Pb content in inoculated roots of Col-0 and *PbE3-2* overexpression plants at 21 days after inoculation with *P. brassicae*. Three plant roots were taken as a mixed sample. n=3 biological replicates. D Col-0 and *PbE3-2* overexpression plants were inoculated with *B. cinerea*, pictures were taken at 24 h post inoculation (hpi). E Disease spot area in C, data are presented as the means ± s.d. (n=10 biological replicates). F Col-0 and PbE3-2 overexpression plants were challenged with *Sclerotinia sclerotiorum*, 30 hpi. G Disease spot area after inoculation of *S. sclerotiorum* in wild-type and *PbE3-2* overexpression plants. Data are presented as the means ± s.d. (n=10 biological replicates). C, E and G were used one-way ANOVA with Kruskal-Wallis test (significance set at *P* ≤ 0.05). Different letters show significant difference. Experiments in a, b and c were independently performed three times with similar results, D, E, F and g with two experiments.

### PbE3-2 promotes the development of clubroot

To characterize PbE3-2 biological functions, we generated stable transgenic *A. thaliana* lines constitutively expressing the PbE3-2 coding sequenceunder the control of the cauliflower mosaic virus 35S (CaMV35S) promoter. Independent homozygous *A. thaliana* lines (OX2, OX5 and OX6) were identified by using western blotting analysis (Appendix Fig. S2). The transgenic plants displayed similar growth phenotypes to Col-0 plants (Appendix Fig. S2). To determine whether PbE3-2 contributes to clubroot disease, transgenic lines were inoculated with *P. brassicae* resting spores, and the occurrence of clubroot disease was investigated at 21 days post inoculation (dpi) (Fig. 2A). The roots of wild-type plants showed the formation of typical galls and correspondingly fewer rootlets, and severe galls (disease classes 5) accounted for 12.8% of the total; besides, the infected plants exhibited purple leaves (Fig. 2A and B). The PbE3-2 transgenic lines showed more severe symptoms than Col-0, with severe galls (disease classes 5) on the roots accounting for 50%~60%, and leaves were dark purple and yellow (Fig. 2 A and B). The disease index of Col-0 was 64.1, while that of transgenic lines was 90.3~91.6 (Fig. 2B). qPCR results revealed that the *P. brassicae* content in the roots of *PbE3-2* transgenic plants was more than 10-fold that of Col-0 (Fig. 2C). These results suggested that *PbE3-2* transgenic plants were more susceptible to *P. brassicae* than Col-0. Although *P brassica* is a biotrophic pathogen, its effector PbE3-2 may also alter the susceptibility of plants to necrotrophic pathogens. Hence, we assessed the susceptibility of *PbE3-2* transgenic lines to the necrotrophic fungi *Botrytis cinerea* and *Sclerotinia sclerotiorum*. As a result, the *PbE3-2* transgenic lines exhibited more significant symptoms of fungal infection than the Col-0 (Fig. 2D-G), indicating that PbE3-2 also modulates the susceptibility of *A. thaliana* to necrotrophic pathogens.

### PbE3-2 suppresses plant immune response

We tested whether PbE3-2 suppress typical immune responses such as reactive oxygen species (ROS) burst, MAPK activation, and immune gene induction in response to PAMP. Chitin-triggered ROS burst was reduced by about 50% in *PbE3-2* transgenic plants relative to Col-0 (Fig. 3A). Moreover, expression level of the *RBOHD* gene was also significantly reduced in *PbE3-2* transgenic plants compared with that in Col-0 (Appendix Fig. S3A). However, a comparative analysis of MAPK activation showed no significant difference between these treatments (Appendix Fig.. S3B). SA is a central component in plant defense against a number of biotrophic pathogens including *P. brassicae* (Volt et al, 2009; Chen et al, 2016). We determined the expression levels of the marker genes in the SA signaling pathway in *PbE3-2* transgenic lines and wild-type plants at 24 h after chitin treatment. The expression of these maker genes was significantly lower compared with that in Col-0 (Fig. 3B and C). These results indicated that PbE3-2 suppresses PTI responses consistently with our observation that PbE3-2 promotes clubroot disease.

**Figure 3.**
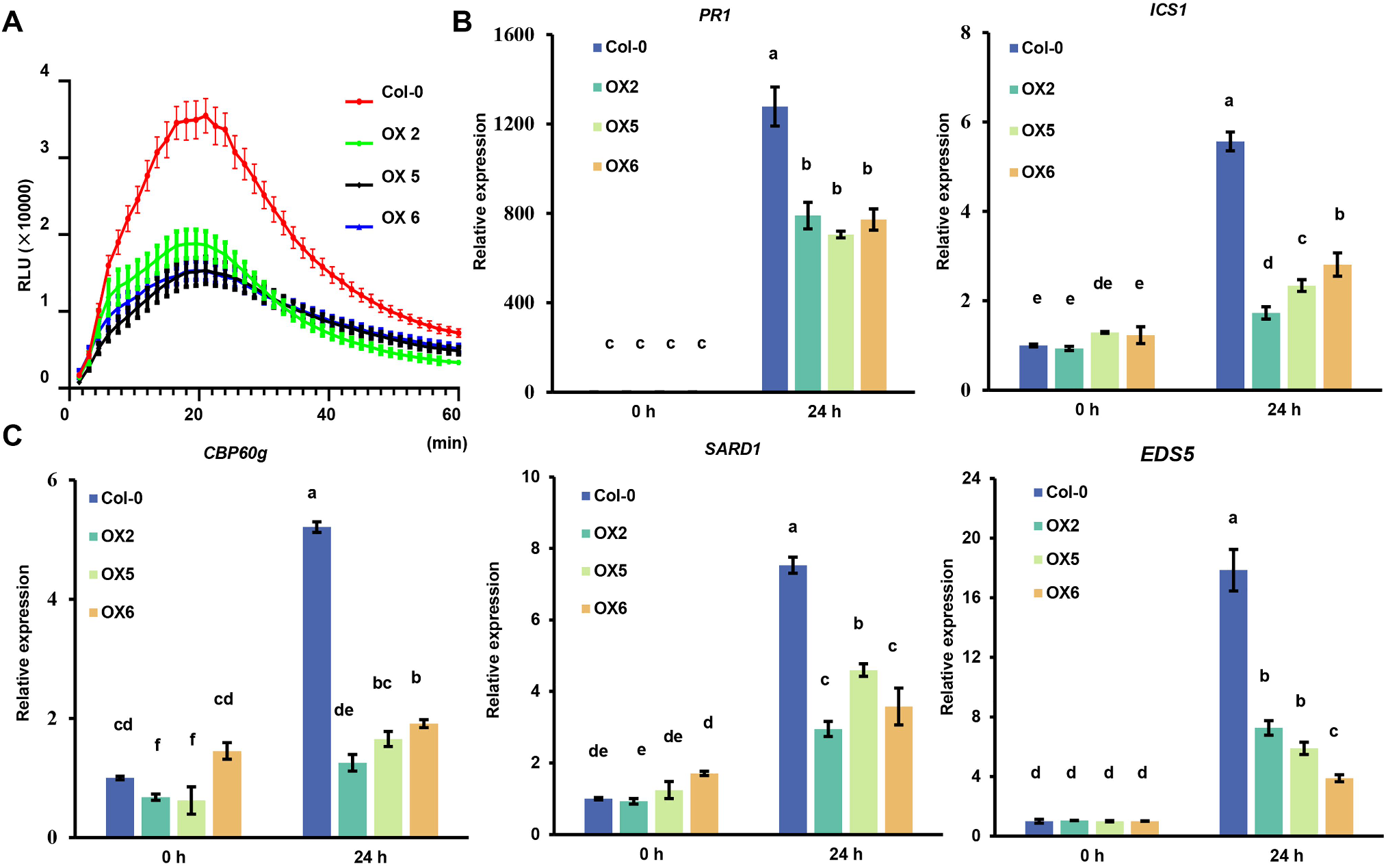
PbE3-2 suppresses plant immune response. A Chitin-induced ROS burst is suppressed in *PbE3-2* overexpression lines. Leave discs from 4-week-old plants were treated with 20 μg/mL chitin and the values represent means ± SE (n=16 biological replicates). B and C Induction of the salicylate pathway genes was compromised in *PbE3-2* overexpression lines. Leaves of 4-week-old plants were infiltrated with 20μg/mL chtitin for 24 h. Data were normalized to ACTIN2 expression in RT-qPCR analysis. Statistical analysis in b and c were performed by one-way ANOVA with Kruskal-Wallis test (significance set at *P* ≤ 0.05). Different letters (a, b and c) are significantly different. n = 3 biological replicates; data are shown as mean ± s.d. All experiments were repeated three times with similar results.

### PbE3-2 interacts with the plant cysteine protease RD21A

To identify the host target of PbE3-2, 35S:PbE3-2-GFP transgenic *A. thaliana* lines infected with *P. brassicae* were prepared. Immunoprecipitation (IP) and liquid chromatography-mass spectrometry (LC-MS) identified 268 potential interacting proteins of PbE3-2 (Table EV2). We next screened *A. thaliana* cysteine protease resistant-to-dehydration 21A (RD21A) in the yeast two-hybrid system (Fig. 4A), and the interaction was validated with a split luciferase complementation assay (Fig. 4B), further examined by Co-IP assay by tagging PbE3-2 with GFP whereas tagging RD21A with FLAG, which were co-expressed in *N. benthamiana* leaves. As shown in Fig. 4C, FLAG-tagged RD21A could be co-immunoprecipitated with GFP-tagged PbE3-2. These results indicated PbE3-2 interacts with RD21A in planta. We further verified the interaction between PbE3-2 and RD21A using an *in vitro* protein-protein pull-down assay. PbE3-2 was fused with His tag and RD21A was fused with glutathione S-transferase (GST), which were purified, incubated, and then immobilized to glutathione sepharose beads. The binding of PbE3-2 to the beads was detected by immunoblotting with the anti-His antibody (Fig. 4D). Together, these results suggested that PbE3-2 directly interacts with RD21A in the plant cell. Subcellular co-localization revealed that PbE3-2-GFP and RD21A-mCherry fluorescence signals were overlapped in the nuclear membrane, plasma membrane, cytoplasm and apoplastic space (Fig. 4E), supporting the notion that PbE3-2 physically interacts with RD21A. Furthermore, BIFC technique also revealed the location of the interaction of PbE3-2 with RD21A (Appendix Fig. S4A), and this result is consistent with the localization of PbE3-2(Appendix Fig. S4B).

**Figure 4.**
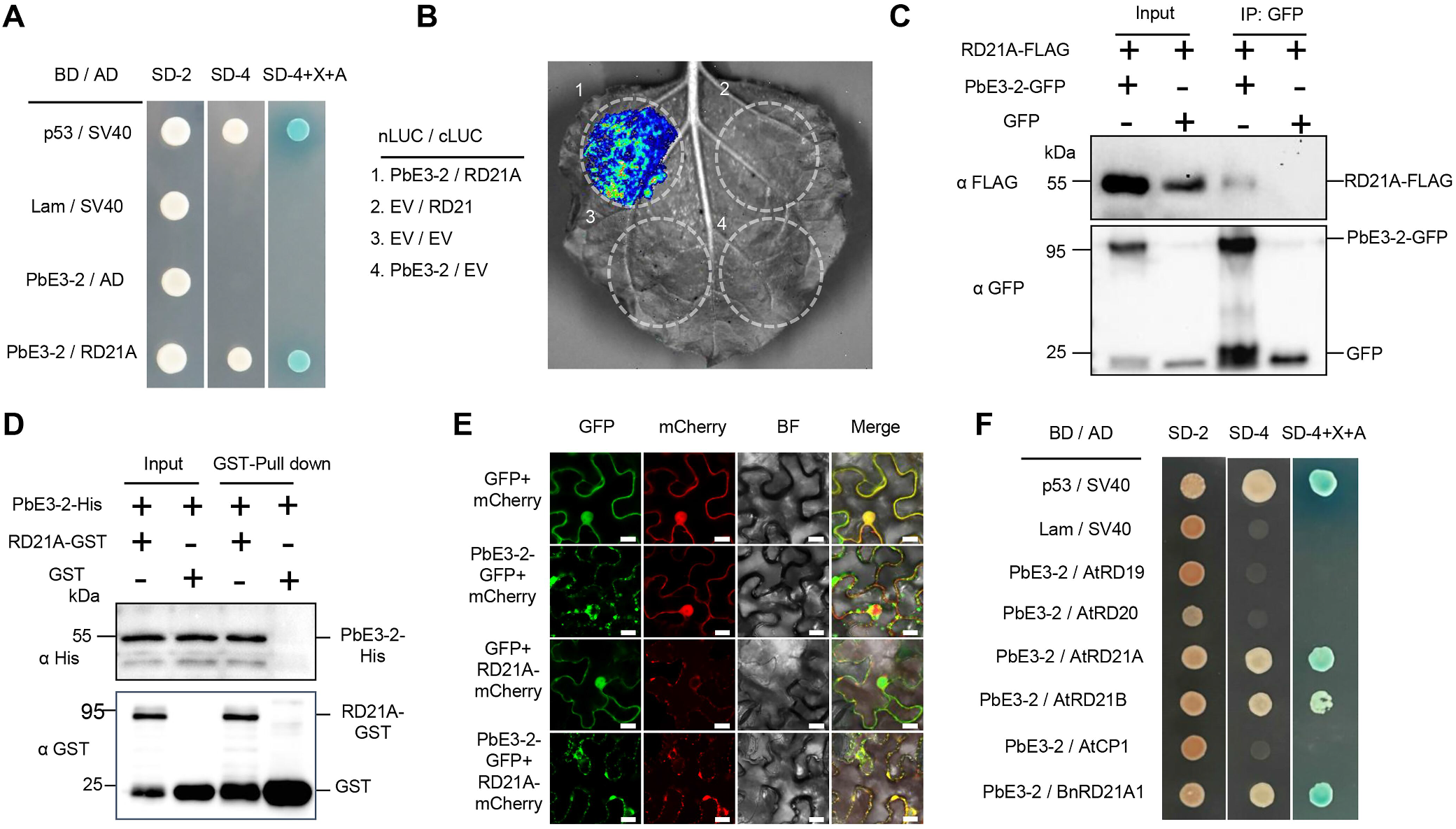
PbE3-2 interacts with *A. thaliana* cysteine protease RD21A. A Yeast two-hybrid (Y2H) assays show that PbE3-2 interacts with RD21A. BD indicated pGBKT7 vector. AD indicated pGADT7 vector. p53+SV40 as positive control, Lam+SV40 as negative control. Yeast cells containing both plasmids were grown on SD-Leu-Trp medium (SD-2) and assessed for interactions on SD-Leu-Trp-His-Ade medium (SD-4) with/without containing X-Gal and aureobasidin A (ABA). B Interaction between PbE3-2 and RD21A in split-LUC assay. PbE3-2-nLUC was co-expressed with RD21A -cLUC in N*. benthamiana* leaves. The luminescence intensity was detected by imaging system within 5 min after the supplement of substrate luciferin. C Co-immunoprecipitation (Co-IP) confirmed that PbE3-2 is associated with RD21A *in vivo*. PbE3-2-GFP or GFP was expressed in *N. benthamiana* together with RD21A-FLAG. Immunoprecipitation was performed with anti-GFP antibody coupled to agarose beads, followed by western blotting. Immunoblots of total proteins (Input) and proteins eluted from the anti-GFP beads (IP) were performed using anti-FLAG and anti-GFP antibodies. D Physical interaction of PbE3-2 and RD21A *in vitro* was verified by GST pull down assay. GST-RD21A was incubated in binding buffer containing glutathione-agarose beads with or without PbE3-2-6 × His, and agarose beads were washed for three times and eluted. Lysis of Escherichia coli (Input) and eluted proteins (Pull-down) from beads were immublotted using anti-HIS and anti-GST antibodies. E Subcellular co-localization of PbE3-2 and RD21A in *N. benthamiana* epidermal cells. Both PbE3-2-GFP and RD21A-mCherry are localized in the cytoplasm and apoplast. The fluorescence of GFP and mCherry was monitored at 2 d post-agroinfiltration using confocal laser scanning microscopy. Bars, 20 μm. F Yeast two-hybrid assays verified the specificity of PbE3-2 interaction with RD21A. PbE3-2 interacted with the Arabidopsis cysteine proteases RD21A and RD21B, but not with the Arabidopsis cysteine proteases RESPONSIVE TO DEHYDRATION 19 (RD19) and cysteine protease 1 (CP1). PbE3-2 also did not interact with RESPONSIVE TO DESSICATION 20. PbE3-2 interacted with RD21 homologs in *Brassica napus* (BnRD21A1, BnaA08g04080D). All experiments were repeated three times with similar results.

To determine the specificity of the interaction between PbE3-2 and RD21A, we cloned the cysteine proteases *AtRD19, AtRD21B* and *cysteine protease 1* (*AtCP1*), RESPONSIVE TO DESSICATION 20 *AtRD20* from *A. thaliana*, and *BnRD21A1* from rapeseed, an ortholog of *A. thaliana* AtRD21A (with an 83.55% similarity at the amino acid level). The results demonstrated that PbE3-2 interacts with AtRD21A, AtRD21B and BnRD21A1, while not with AtRD19, AtRD20 and AtCP1 in yeast cells (Fig. 4F). These results suggested that PbE3-2 interacts explicitly with cysteine proteases in the RD21 clade. Three orthologs of AtRD21A in rapeseed that can interact with PbE3-2 were examined by split luciferase complementation assays (Appendix Fig. S5). The results demonstrated that the interactions between PbE3-2 and RD21 orthologs are conserved.

Full-length AtRD21A contains a signal peptide, an auto-inhibitory precursor protein, a protease domain, a proline-rich region, and a granule protein domain (Yamada *et al*, 2001; Gu *et al*, 2012; Pogorelko *et al*, 2019). According to available information, this precursor protein has undergone extensive post-translational processing, resulting in the forms of intermediate RD21A (iRD21A, removal of signal peptide and autoinhibitory pro-domain) and mature/active RD21A (mRD21A, consisting of only the protease domain). To characterize the structural requirements of the interaction between RD21A and PbE3-2, RD21A and its four truncated derivatives were fused to the GAL4 activation domain and analyzed for potential interactions with PbE3-2 (Appendix Fig. S6A-C). PbE3-2 interacted with RD21A, iRD21A, mRD21A (Appendix Fig. S5B). Among the four constructs, the one containing the protease domain exhibited a stronger interaction with PbE3-2, suggesting that the protease domain of RD21A is responsible for its interaction with PbE3-2. Besides, the granule protein domain had a much weaker interaction with PbE3-2 in the colony growth assay (Appendix Fig. S5C), and thus is not essential to the interaction with PbE3-2. Split luciferase complementation imaging assays were performed to further verify the interactions between RD21A constructs and PbE3-2. All three forms of RD21A exhibited fluorescence (Appendix Fig. S6D), indicating their interactions with PbE3-2 *in vivo*.

### PbE3-2 ligase ubiquitinates RD21A

As PbE3-2 is a E3 ubiquitin ligase and interacts with RD21A, we investigated whether RD21A is a substrate of PbE3-2. We reconstructed the Arabidopsis ubiquitination cascade in vitro as previously reported (Han *et al*, 2017). We incubated recombinant AtUBA1-S (E1), PbE3-2-Myc, RD21A-HA, AtUBC8-S (E2), and His-FLAG-UBQ10 (ubiquitin) and tested for ubiquitination. The results showed that RD21A had a laddering pattern, and its ubiquitination was only observed when all components were present (Fig. 5A). Besides, the PbE3-2-C423S mutant showed no self-ubiquitination, and could not ubiquitinate RD21A (Fig. 5A), indicating that the cysteine residue at position 423 in the RING domain is essential for RD21A ubiquitination. Interestingly, the mutation (C423S) did not affect the interaction between PbE3-2 and RD21A in yeast two-hybrid assays (Fig. 5D), suggesting that the E3 ubiquitin ligase activity of PbE3-2 is not required for the physical interaction between the two proteins. We then set out to determine whether other two forms of RD21A can be ubiquitinated by PbE3-2. The results showed that all of the three forms of RD21A, including full length RD21A, iRD21A and mRD21A, were ubiquitinated by PbE3-2 (Fig. 5B). We next tested whether PbE3-2 can promote the degradation of RD21A *in vivo*. FLAG-tagged RD21A was transiently co-expressed with PbE3-2-GFP or PbE3-2-C423S-GFP in *N. benthamiana* leaves, and the accumulation of RD21A was monitored by western blot. As shown in Fig 5E, RD21A was accumulated to a relatively higher level when co-expressed with PbE3-2-C423S in comparison with PbE3-2. Application of the 26S proteasome degradation inhibitor MG132 significantly inhibited the degradation of RD21A (Fig. 5E). These results suggested that PbE3-2 directly interacts with RD21A and promotes its degradation through the 26S proteasome pathway.

**Figure 5.**
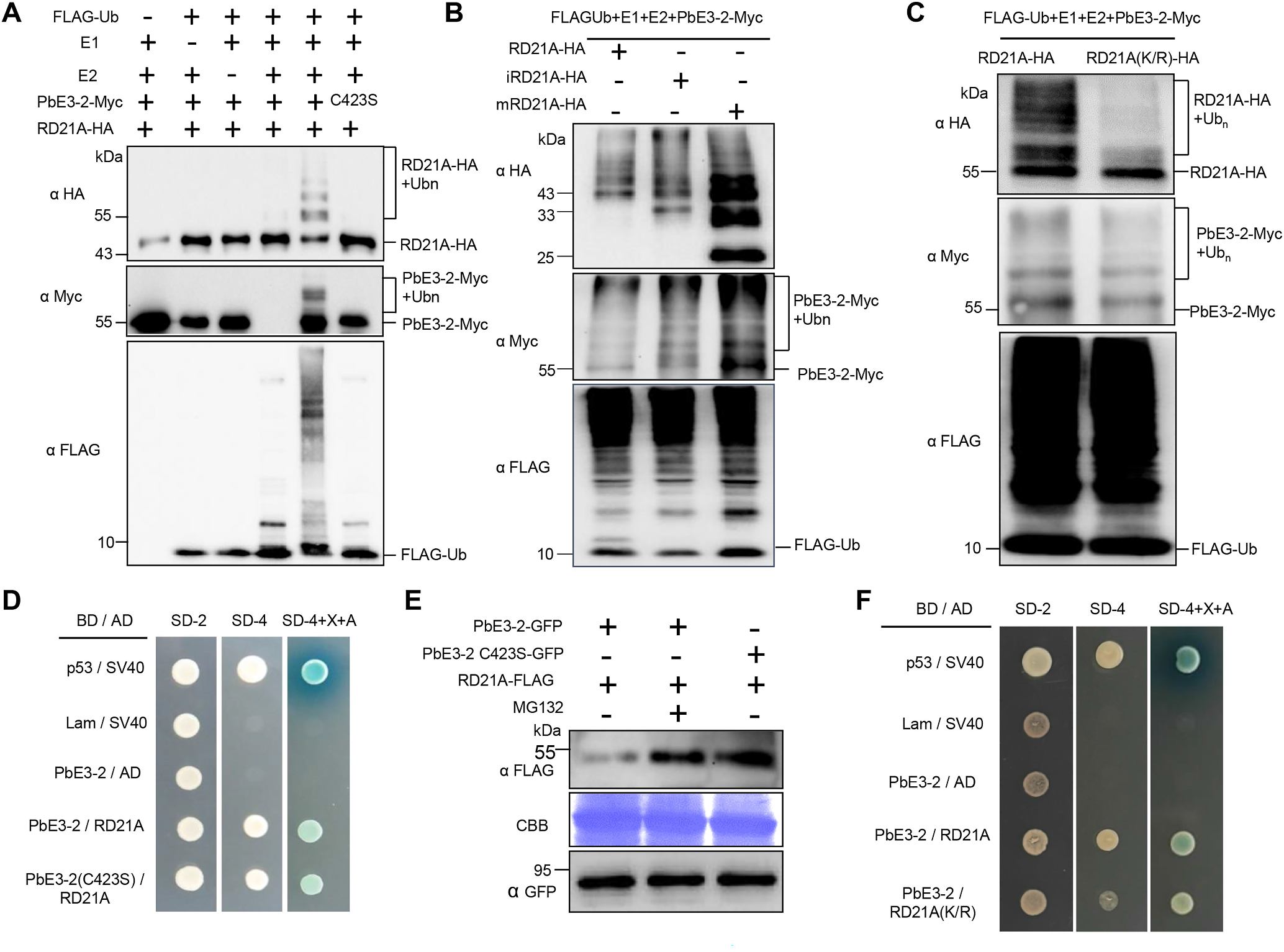
RD21A is a ubiquitinated substrate of PbE3-2 and is mediated by degradation through the 26S proteasome. A PbE3-2 ubiquitinates RD21A whereas PbE3-2(C423S), an enzyme active site mutant of PbE3-2, fails to ubiquitinate RD21A. B Yeast two-hybrid assay shows that PbE3-2(C423S), an enzyme active site mutant of PbE3-2, interacts with RD21A. C RD21A, iRD21A and mRD21A are ubiquitinated by PbE3-2. D RD21A was degraded by PbE3-2. PbE3-2 or PbE3-2 C423S was co-expressed with RD21A in *N. benthamiana* leaves. The amount of RD21A was detected by immunoblotting with an a-FLAG antibody (top). Protein loading was shown by Coomassie blue staining for RBC (bottom). E RD21A (K/R), a ubiquitination site mutant of RD21A, is ubiquitinated by PbE3-2 at a much lower level. F Yeast two-hybrid assay showing that RD21A(K/R), a ubiquitination site mutant of RD21A, interacts with PbE3-2. All experiments were repeated three times with similar results.

### RD21A ubiquitination sites

Since RD21A is ubiquitinated by PbE3-2, we predicted and identified the ubiquitination sites in RD21A protein sequences. We purified the ubiquitin-conjugated RD21A protein and identified the ubiquitination sites by mass spectrometry analysis, and identified 13 lysine sites (Table EV3). We then generated a dominant-negative ubiquitin mutant RD21A (K/R) with the substitution of the 13 predicted Lys (K) residues by Arg (R) (Fig. 5C). As a result, the ubiquitination degree in RD21A (K/R) was significantly reduced compared with that in the wild type RD21A. However, the interaction between PbE3-2 and RD21A (K/R) in yeast cells was not affected (Fig 5F). These results suggested that the 13 Lys residues in RD21A play an essential role in ubiquitination.

### RD21A mediates plant disease resistance

To test whether RD21A contributes to resistance against *P. brassicae*, Col-0 and *rd21a* mutant plants were inoculated with *P. brassicae* for 21 days. The non-inoculated *rd21a* mutant showed no observable change in morphology compared with Col-0 (Appendix Fig. S3). However, the inoculated *rd21a* mutant exhibited much more severe symptoms than Col-0 with yellow leaves and severe galling, while the control plants showed galls with lateral roots (Fig. 6A). The percentage of severe galls (disease levels of 4 and 5) was 53% in Col-0 and 91% in *rd21a*. The disease index was 67.2 for Col-0, while 87.0 for *rd21a* (Fig. 6B). *P. brassicae* content in the galls of Col-0 was significantly lower than that in the *rd21a* mutant (Fig. 6C). These results indicated that *rd21a* was more susceptible to *P. brassicae* than Col-0. Besides, *rd21a* was more susceptible to *B. cinerea* than Col-0 (Fig. 6D). Since it was already known that PbE3-2 suppresses host defense response (Fig. 3), we tested whether RD21A mediates the defense response by comparing the ROS production and transcript levels of defense marker genes (Fig. 6E-F). Comparative analysis of chitin-triggered ROS burst and SA signaling pathway marker gene expression revealed that they were significantly reduced in the *rd21a* mutant than in Col-0 (Fig. 6E-F). Taken together, the *PbE3-2* transgenic plants and *rd21a* mutant had consistent disease susceptible phenotype and suppression of defense response, suggesting that PbE3-2 alters host defense responses by degrading plant RD21A protein through the 26S proteasome pathway, thereby promoting pathogen infection.

**Figure 6.**
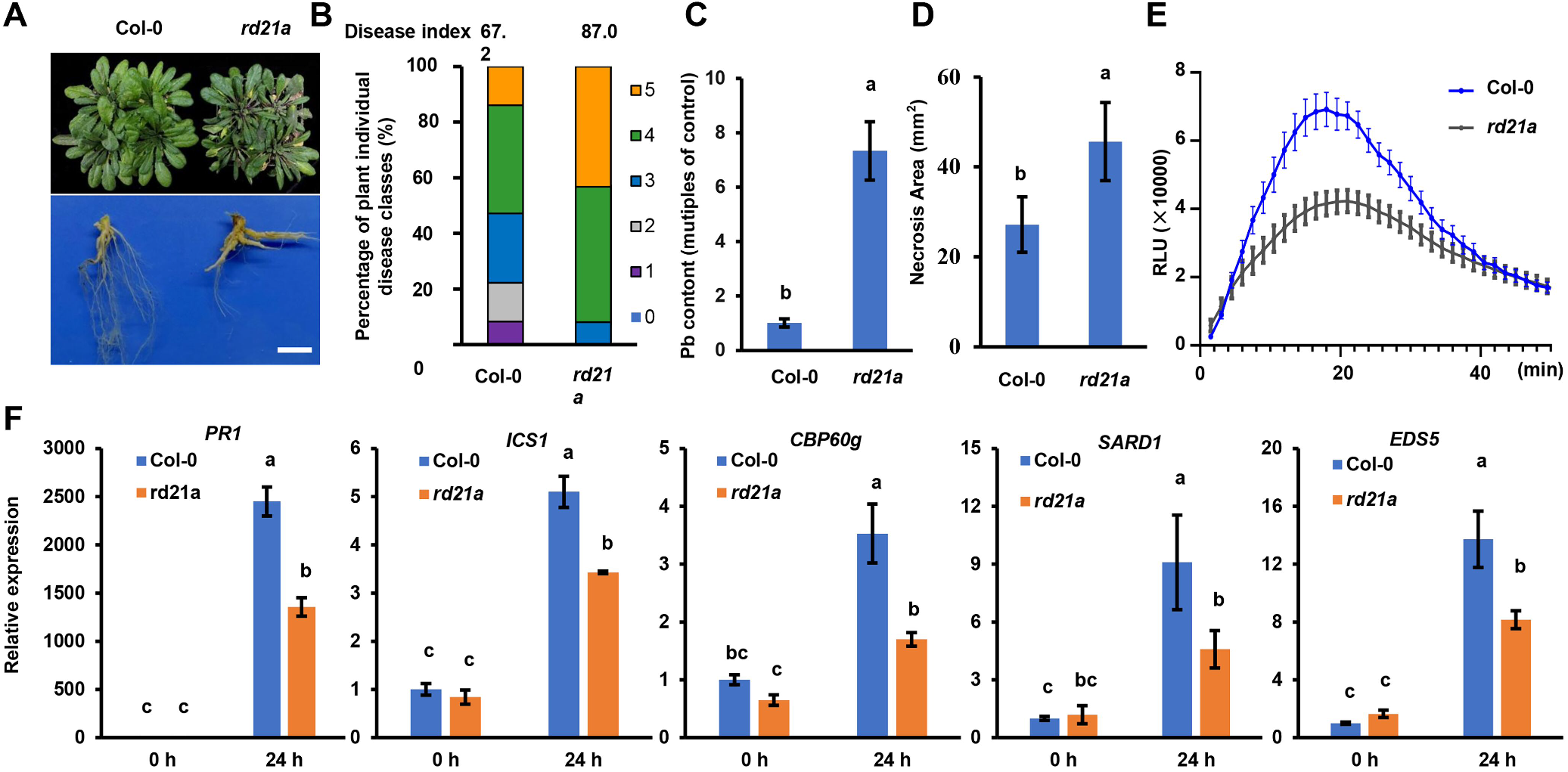
*rd21a* mutants exhibit susceptibility to pathogens and reduced levels of immunity. A Phenotype of Col-0 and *rd21a* at 21 days after *P. brassicae* inoculation. B Disease index of Col-0 and *rd21a*. The percentages of plants in the individual disease classes are shown, Col-0 and *rd21a* used 35-40 plants for analysis. C Pathogen DNA quantification (Pb) by quantitative PCR, which is expressed as the percentage of the mean Pb content in inoculated roots of Col-0 and *rd21a* at 21 days after inoculation with *P. brassicae*. Three plant roots were taken as a mixed sample, n = 3 biological replicates. Significant difference was determined by Student’s t-test (two-tailed) Different letters (a and b) are significantly difference. Data are shown as mean ± s.d. D Disease spot area after inoculation of *B. cinerea* in wild-type and *rd21a* at 24 h. Significant difference was determined by Student’s t-test (two-tailed) Different letters (a and b) are significantly difference. Data are shown as mean ± s.d. E Chitin-induced ROS burst is suppressed in *rd21a* mutant. Leave discs from 4-week-old plants were treated with 20 μg/mL chitin and the values represent means ± SE (n =16 biological replicates). F Induction of salicylate pathway genes was compromised in *rd21a* mutant. Leaves of 4-week-old plants were infiltrated with 20 μg/mL chitin for 24 h. Data were normalized to ACTIN2 expression in RT-qPCR analysis. Statistical analysis was performed by one-way ANOVA with Kruskal-Wallis test (significance set at *P* ≤ 0.05). Different letters (a, b and c) are significantly different. n = 3 biological replicates; data are shown as mean ± s.d. All experiments were repeated three times with similar results.

### Role of PbE3s in the pathogenesis of *P. brassicae*

We next tested whether PbE3-3 and PbE3-4 perform the same function as PbE3-2. As a result, both of them were capable of self-ubiquitination (Appendix Fig. S7A). In addition, PbE3-3 and PbE3-4 can suppress the BAX-triggered cell death in *N. benthamiana* (Appendix Fig. S7B), suggesting that PbE3-3 and PbE3-4 can suppress plant immunity during the infection of host plants. In addition, PbE3-3 and PbE3-4 interact with RD21A in a yeast two-hybrid assay (Appendix Fig. S7C). Interestingly, in the reconstituted ubiquitination system in *E. coli* cells, PbE3-3 and PbE3-4 also ubiquitinated RD21A (Appendix Fig. S7D). RD21B is a homologue of RD21A with a 78.9% similarity at the amino acid level (Appendix Fig. S8A) and also interacts with PbE3-2 (Fig 4F), therefore they may be functionally redundant. To confirm this possibility, we determined whether RD21B is the substrate of these E3 ubiquitin ligases. As expected, we observed the ubiquitination of RD21B by PbE3-2 (Appendix Fig. S8B).

## Discussion

The rapeseed and cruciferous vegetable industry is facing unprecedented challenges from *P. brassicae*. In-depth understanding of the pathogenesis of *P. brassicae* is urgent to develop effective management strategies. In this study, we identified the secreted E3 ubiquitin ligase PbE3-2 of *P. brassicae*, which takes the cruciferous cysteine protease RD21 as a virulence target and mediates its degradation (Fig. 7), thus providing new insights into the pathogenesis of *P. brassicae*.

**Figure 7.**
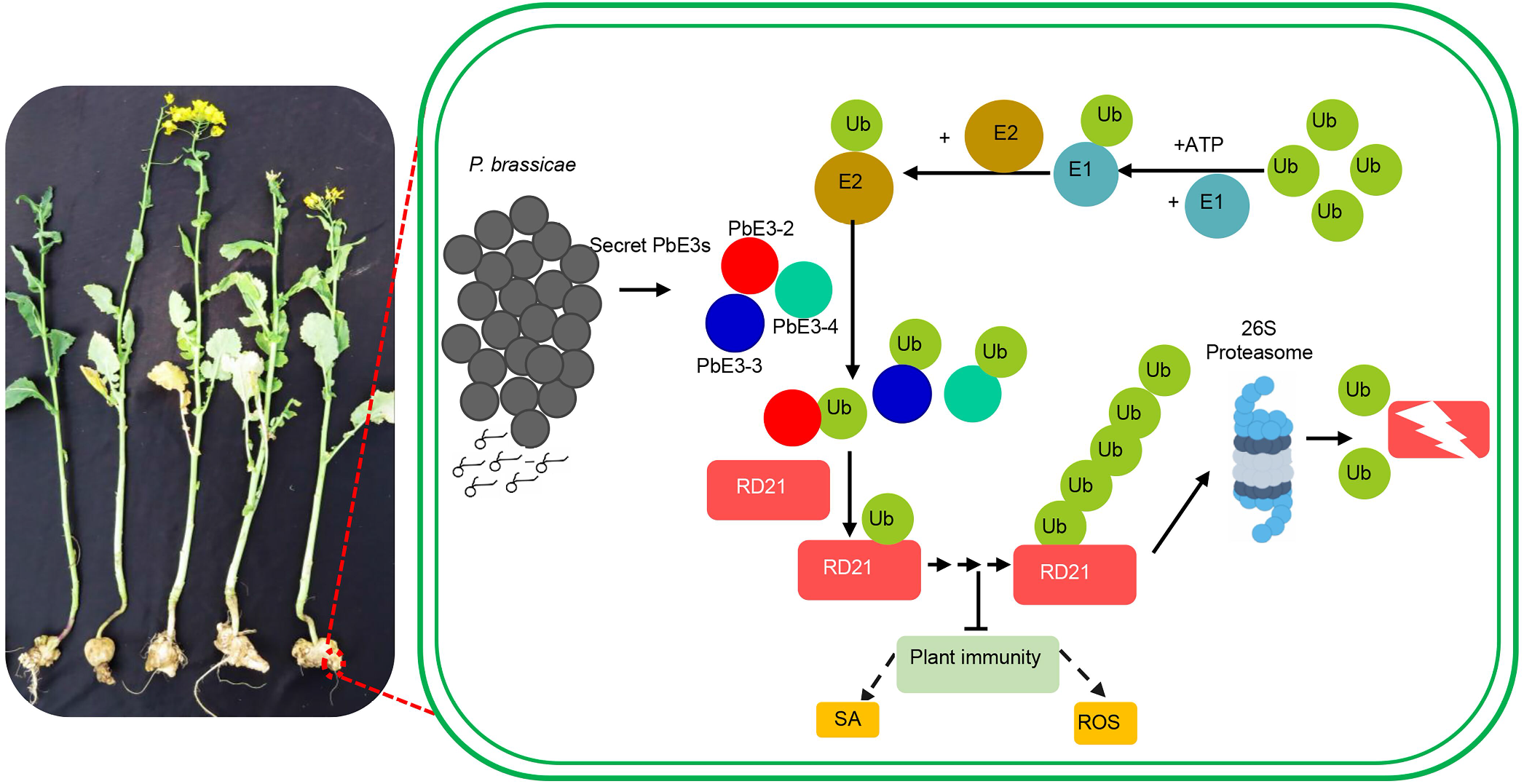
Working model of the mechanism for PbE3 to regulate plant immunity. The E3 ubiquitin ligase secreted by *P. brassicae* directly targets the cysteine protein RD21 of the host plant, leading to its degradation via the ubiquitin-proteasome pathway. Degradation of RD21, a positive regulator of plant immunity, leading to a reduction in the level of plant immunity and promoting infestation of *P. brassicae*.

In the present study, we identified PbE3-2, PbE3-3 and PbE3-4 are secreted E3 ubiquitin ligase in the *P. brassicae* (Fig. 1, Appendix Fig. S7 and Table EV1). Furthermore, overexpression of *PbE3-2* in *A. thaliana* resulted in significant susceptibility to *P. brassicae* (Fig. 2), suggested that secreted E3 ubiquitin ligases play important roles in clubroot development. It has been reported that some pathogens secret effector to target host ubiquitin systems to inhibits plant immunity to establish successful infection (Marino *et al*, 2012; Park *et al*, 2012; Ashida & Sasakawa, 2017). While in this mysterious protist, they directly secreted E3 ubiquitin ligase, it would be interesting to determine the targets of these secreted E3 ubiquitin ligases, and would potentially expand our knowledge of the repertoire of pathogenesis.

To explore how PbE3-2 regulated plant immunity, we used IP and LC-MS to screen the candidate interacting proteins. Through yeast two-hybrid assay, split luciferase complementation, Co-IP, pull down and co-localization assays, we demonstrated that PbE3-2 interacts with the *A. thaliana* cysteine protease RD21A both *in vivo* and *in vitro* (Fig. 4). *RD21A* has been identified as one of the drought-induced marker genes and is well documented (Koizumi *et al*, 1993). Several RD21A interact proteins have been identified that may help us to understand the biological and biochemical roles of RD21A in plants (Davies *et al*, 2015; Rustgi *et al*, 2017), RD21A is not only involved in plant immunity but also plays a key role in plant development (Rustgi *et al*, 2017). RD21A is involved in the resistance of Arabidopsis to bacteria, fungi, as well as nematodes, and that mutating *RD21A* increases plant susceptibility (Shabab *et al*, 2008; Lozano-Torres *et al*, 2012; Shindo *et al*, 2012; Pogorelko *et al*, 2019; Liu *et al*, 2020). Proper RD21A homeostasis is critical as excessive accumulation of RD21A can lead to autoimmunity and cell death, while insufficient accumulation of RD21A may result in susceptibility to pathogens (Lampl et al, 2013; Boex-Fontvieille et al, 2015). Therefore, the level of RD21A in plant cells is tightly regulated by various biochemical reactions. In this study, we confirmed that knocking out *rd21a* is more susceptible to *P. brassicae* (Fig. 6). ROS production and transcript levels of defense marker genes in the chitin-triggered immune responses in the *rd21a* plants were decreased significantly than that of Col-0 (Fig. 6). RD21A can be ubiquitinated by PbE3-2, PbE3-3 and PbE3-4 *in vitro* (Fig. 5, Appendix Fig. S7), RD21B, a homologue of RD21A, can be ubiquitinated by PbE3-2 (Appendix Fig. S7), all of these suggest ubiquitination of RD21 by secreted E3 ubiquitin ligase in the *P. brassicae*. In previous reports, RD21A level is also regulated by the plant ubiquitination system (Kim & Kim, 2013). RD21A can interact with the ubiquitin E3 ligase SINAT4 to participate in drought-induced immunity (Liu *et al*, 2020). Unlike previously reported, we identified pathogen secreted ubiquitin E3 ligase to regulation RD21A level.

The papain-like cysteine proteases (PLCPs) have important, complex and not fully understood functions for the regulation of plant immunity. PLCPs contribute to plant resistance to a wide range of pathogens. Tomato immune proteases RCR3 and PIP1 contribute to defense against fungal and oomycete pathogen (Rooney *et al*, 2005; Kaschani *et al*, 2010; Paulus *et al*, 2010). Maize immune signalling peptide 1 (Zip1) was released by PLCP and can strongly trigger the accumulation of SA in leaves (Ziemann *et al*, 2018). In addition, protease could cleave microbial peptides to elicit defense responses (Wang *et al*, 2021). Since PLCP plays an important role in plant immunity, many pathogens have evolved effector proteins that target PLCP. Huanglongbing-associated pathogen secreting SDE1 directly interacts with citrus PLCP and inhibits its protease activity (Clark *et al*, 2018). *Pseudomonas syringae* secretes CIP1 to inhibit tomato immune-associated protease thereby promoting virulence (Shindo *et al*, 2016). The cyst nematode effector Hs4E02 targets plant defense protease and achieves virulence by altering its subcellular localization (Pogorelko *et al*, 2019). In the present study, we identified *P. brassicae* directly targets the plant defense-associated cysteine protease RD21 to promote virulence, which increases our understanding of the pathogenesis of protist. Therefore, in the future, it will be interesting to investigate how RD21A regulated plant immunity in the clubroot development.

Taken together, our findings demonstrate that cysteine proteases RD21 are a complex component of plant immunity, can be degraded directly by pathogen as a ubiquitinated substrate. These results provide a basis for understanding the pathogenesis of *P. brassicae* and developing germplasm resources for *P. brassicae* resistance through genetic manipulation.

## Materials and Methods

### Pathogen Strains and Inoculation

*P. brassicae* strain ZJ-1 was originally isolated from a diseased plant in a rapeseed field in Zhijiang County, Hubei Province, China (Chen *et al*, 2016). Resting spores of *P. brassicae* were extracted from the galls of rapeseed and stored at 4°C. Fourteen-day-old *A. thaliana* seedlings were inoculated with 1 mL of the resting spore suspension (1.0 × 10^7^ spores/mL) through the soil around each plant, and the phenotype was investigated at 21 days post inoculation (dpi). The disease severity was assessed using a scoring system of 0–5 modified from a previous study (Siemens *et al*, 2002). 0, no disease; 1, very small galls mainly on lateral roots, no damage on main roots; 2, small galls covering the main roots and a few lateral roots; 3, medium to large galls on the main roots; 4, severe galls on lateral roots, main roots or rosettes, with complete destruction of fine roots; 5, completely expansion and rotting of roots.

### Plants and growth conditions

The Arabidopsis and *Nicotiana benthamiana* plants were grown in a growth chamber at 12 h day, 12 h night, 22°C and 60% relative humidity. CRISPR/Cas9 approach was used to introduce a mutation into the N-terminus of RD21A in the Col-0 background to obtain the *rd21a* mutant (Liu *et al*, 2020). *35S:PbE3-2* transgenic *A. thaliana* line was generated by using Agrobacterium-mediated transformation by floral dipping (Clough & Bent, 1998).

### Extraction of apoplastic fluid

Extraction of plasmatic extracellular fluid was conducted in reference to the previous protocol (O’Leary et al, 2014). Sterile water was infiltrated by vacuum pump into the leaves of *N. benthamiana* with transient expression of PbE3-2 or PbE3-2^ΔSP^ protein. The apoplast washing fluid was recovered by centrifugation and concentrated for western blotting to verify that proteins were secreted into the apoplast.

### Ubiquitination assay

Ubiquitination assay were performed by reconstituted ubiquitination reactions in *Escherichia coli* (Han et al, 2017). Arabidopsis ubiquitin UBQ10 was constructed into the prokaryotic expression vector pET28a. AtUBA1 and RD21A were constructed into the compatible dual expression vector pCDFDuet, and AtUBC8 and PbE3-2 were constructed into the compatible dual expression vector pACYDuet. Strains lacking any of these components were used as negative controls. Self-ubiquitination reactions were performed with strains lacking the substrate. After induction with 0.5 mM β-D-1-thiogalactopyranoside (IPTG) at 20°C for 18 h, the purified proteins from the crushed organisms were subjected to western blot for each fraction to determine the occurrence of ubiquitination.

### Quantification of *P. brassicae* DNA content in infected roots

DNA was extracted from root samples using the cetyltrimethylammonium bromide (CTAB) method (Allen et al, 2006). Quantitative PCR was performed using iTaq Universal SYBR Green supermix (BioRad) on a CFX96 real-time PCR system (BioRad) as described by Chen et al (2016). Arabidopsis actin gene AT3G18780 was used as an internal control for data normalization. The qPCR primers used are listed in Table EV4.

### ROS assay and MAPK assay

ROS assay and MAPK assay was performed as previously described (Qi et al, 2022). Leaves discs from 4-week-old Arabidopsis plants were incubated in 96-well plates with 100 μL H_2_O overnight. 100 μL reaction solution containing 5 μM L-012 (Wako,120-04891, Japan), 10 μg/mL horseradish peroxidase (Sigma, P6782, USA) and 20 μg/mL chitin were added to 96-well plates.Using a Multimode Reader Platform (Tecan Austria GmbH, SPARK 10M) with a reading interval of 1.5 min with 50 min per treatment to measure ROS production. Five leaves discs were transferred to 6-well plates with H2O overnight, and then treated with 20 μg/mL chitin for 5, 15 or 30 min, respectively. Extracted protein and used α-pErk1/2 antibody (CST, 9101, USA, 1:2000) and secondary antibody goat anti-rabbit IgG-HRP (Abclonal, AS014, China, 1:10000) to detect the level of pMPK3, pMPK4, and pMPK6.

### Immunoprecipitation (IP) and liquid chromatography–mass spectrometry (LC-MS) analysis

The IP experiment was performed as previously described (Yang et al, 2018). Using IP buffer (Beyotime, P0013, China) to extract the total protein from *PbE3-2-GFP* transgenic plants. About 50 mL of protein extract was incubated with 100 μL of anti-GFP agarose beads (chromotek, gta-20, Germany) for 12 h at 4°C on a rotary shaker. The beads were then collected by centrifugation at 1000 g for 1 min at 4°C and then washed 3 times with 1 ml of IP buffer. The bound proteins were eluted from the beads by boiling for 5 min. The proteins were sent to HOOGEN BIOTECH company (Shanghai, China) for LC-MS analysis (Q Exactive, Thermo Fisher).

### Yeast two-hybrid system

The cDNA encoding the PbE3-2 without signal peptide was cloned into pGBKT7 vector as bait, and the candidate proteins identified from LC-MS were constructed into pGADT7 as prey. Yeast two-hybrid gold yeast cells were transformed with bait and candidate prey plasmids using polyethylene glycol/LiAc-mediated yeast transformation. The transformation mix was spread on SD medium-Leu-Trp(SD-2, selects for both plasmids) and positive clones were further confirmed on SD medium-Trp-Leu-His-Ade (SD-4) containing X-α-Gal and Aureobasidin A (AbA).

### Co-IP assay

Plasmids 35S:RD21A-FLAG together with 35S:PbE3-2-GFP or 35S: GFP were transiently expressed in *N. benthamiana*. 2 days after *Agrobacterium* infiltration, total proteins were extract with IP buffer, then immunoprecipitation was performed with α-GFP agarose (chromotek, gta-20, Germany) at 4°C for 1 h, the agarose beads were collected and washed with IP buffer for 3 times and immunoblot with an anti-FLAG antibody.

### Split-luciferase complementation assay

The split-luciferase reconstitution assay was performed as described previously (Chen et al, 2008). *Agrobacterium* carrying the specified nLUC and cLUC constructs were co-infiltrated into 4-week-old *N. benthamiana* leaves. About 1 mM luciferin was infiltrated 48 h after infiltration and luminescence was imaged within 10 min.

### Statistical analysis

Statistical significance was determined by conducting Student’s t tests or one-way ANOVA using GraphPad Prism 8.0.

## Data availability

The data are available on request from the corresponding author.

## Acknowledgements

We thank Dr. Yi Liu from Chinese Academy of Sciences for the seeds of *rd21a* mutant. We thank Dr. Dongping Lu from Institute of Genetics and Developmental Biology for sharing the in *vitro* ubiquitination assay system. We thank Dr. Yangrong Cao from Huazhong Agricultural University for sharing the split-LUC vectors. We thank Dr. Kenichi Tsuda, Shengyang He and Kabin Xie for critical reading of the manuscript and constructive suggestions. We thank research associates at Center for Protein Research (CPR), Huazhong Agricultural University, for technical support. We would like to thank the State Key Laboratory of Agricultural Microbiology Core Facility for assistance in confocal microscopy. This research was financially supported by the National Natural Science Foundation of China (32172371), the Fundamental Research Funds for the Central Universities (2662020ZKPY003), the earmarked fund of China Agriculture Research System (CARS-13), and collaborative Fund of Huazhong Agricultural University and Agricultural Genomics Institute at Shenzhen (SZYJY2021007).

## Author contributions

C.L. and T.C. conceived the project and designed experiments. C.L. performed most of the experiments. C.L., S.F.L., L.F. and Q.Q.W. performed experiments and analyzed the data. J.S.C., J.T.X., Y.L., Y.P.F. and D.H.J. analyzed the data and provided advices that helped shape the research. C.L. and T.C. wrote the manuscript with input from all co-authors.

## Supporting Information

**Appendix Fig. S1.** Analysis of secreted PbE3s.

**Appendix Fig. S2.** Growth phenotype of Col-0, *PbE3-2* overexpression plants and *rd21a* mutant.

**Appendix Fig. S3.** Chitin triggered *RBOHD* gene expression and MAPK activation in PbE3-2 overexpression lines.

**Appendix Fig. S4.** Bimolecular fluorescence complementation (BiFC) on the interaction between PbE3-2 with RD21A in planta and subcellular localization of PbE3-2.

**Appendix Fig. S5.** PbE3-2 interacts with *B. napus* BnRD21A.

**Appendix Fig. S6.** PbE3-2 interacts with truncated RD21A.

**Appendix Fig. S7.** PbE3-2, PbE3-3 and PbE3-4 interacted with RD21, respectively.

**Appendix Fig. S8.** PbE3-2 ubiquitinate RD21A and RD21B, respectively.

**Table EV1.** The information of E3 ubiquitin ligases in the genome *of P. brassicae* ZJ-1.

**Table EV2.** List of potential PbE3-2 interacting partners identified by IP-MS/MS analysis.

**Table EV3.** Ubiquitination sites by LC-MS.

**Table EV4.** Primers used in the study.

